# A Generative Virtual Tissue Model Enables Computational Design of Therapeutic Perturbation Strategies

**DOI:** 10.64898/2026.08.12.743536

**Authors:** Yunrui Lu, Willis Zhang, Yu-Jen Chen, Jiayi Yin, Lydia Chen, Kevin Fleisher, James Gornet, Rex Liu, Zitong Jerry Wang, Yan Poon, Yuning You, Matt Thomson

## Abstract

Computational design has transformed many fields of engineering, where simulators can explore millions of candidate design configurations before experimental development and testing. Therapeutic design in biomedicine has resisted computational design approaches because disease progression and therapeutic response emerge from interactions among many cell types within human tissue, governed by biochemical parameters that are largely unknown and potentially unknowable. Here, we introduce the Cell Interaction Foundation Model (CIFM), a virtual tissue model that forward-simulates the transcriptional dynamics of cells in human tissue under arbitrary therapeutic conditions based upon a spatial transcriptomic seed. CIFM is a geometric graph neural network trained by self-supervised masked-transcriptome prediction on millions of cellular microenvironments spanning human tissue types and disease states; generative, auto-regressive, monte-carlo play-out, then, simulates transcriptional dynamics under combinatorial perturbations from a spatial transcriptomic seed. We validate CIFM by showing accuracy gains in gene expression prediction and imputation, disease classification, recapitulation of perturbation responses in prostate cancer models, and recovery of T cell-tumor signaling measured in cell–cell sequencing experiments. Beyond such conventional tasks, CIFM enables target identification and therapeutic design through generative tissue simulation play-outs. Analyzing over 10^6^ single and combinatorial perturbations, CIFM designs immunotherapy strategies for cancer and autoimmune disease that exploit combinatorial manipulation of signaling pathways to induce or suppress immune activation. Broadly, CIFM shows how generative artificial intelligence methods can be applied to model emergent behavior in highly interacting biological systems, yielding new approaches to fundamental understanding of tissue behavior as well as large-scale therapeutic design.

## Introduction

A human tissue is a distributed computational system[1, 2, 3], and the therapeutic programming of tissue behavior requires methods that predict how computations within cells and communication between cells generates tissue states in health and disease. In a tissue each cell senses its local environment and computes a gene expression program[4] while exchanging hundreds of different signaling molecules with neighboring cells. Thus, the physiological state of a tissue, inflamed or suppressed, healthy or diseased, emerges from the collective behavior of thousands of tissue-resident and immune cells interacting through a large and heterogeneous repertoire of signaling molecules and pathways[5, 6, 7, 8].Reprogramming collective tissue states is increasingly the central problem in medicine, because the most promising therapies in cancer and autoimmunity, like checkpoint inhibitors, act not on a single cell type but on interactions between tissue-resident cells and immune cells[6, 9]. Therapeutic failures are correspondingly often driven by unpredicted emergent dynamics[10]. Checkpoint inhibitors fail when signaling networks between cell types, for example between macrophages and tumor cells, exclude immune cells from a tumor, producing immunosuppression[9], or when long-range positive feedback loops drive uncontrolled immune cell recruitment and hyper-inflammation, i.e. cytokine storm[11]. Forward prediction of cellular dynamics at tissue scale could enable large scale design of new therapeutic interventions, adjuvants matched to specific patients and microenvironmental configurations, and a deeper understanding of how function in biology emerges through interactions between cellular components[12, 13].

Engineering disciplines confront emergence during design through mathematical and computational models that predict how interactions between individual components produce new behaviors at larger length and time scales, but biomedicine has no comparable approach for therapeutic design and tissue programming. Models of electronic circuits, materials, and airflow allow millions of configurations to be explored[14], so that designs beyond the reach of intuition are found by search rather than intuition[15, 16]. Tissue state has resisted forward modeling because of the intricate chemical complexity of cellular regulatory networks. To account for the molecular complexity of chemical interactions within a tissue, a bottom-up mechanistic model would require identification of billions of biochemical parameters that cannot be measured systematically at tissue scale[17, 18], while the reductionist alternative of tracking a handful of pathways and cell types discards precisely the multi-cell couplings that drive therapeutic failure[19]. Virtual cell models attempt to predict the behavior of cells in isolation[4, 20, 21, 22], leaving emergent tissue dynamics as a future challenge. Current models that operate at tissue scale, meanwhile, predict cell-population composition[13] or classify pathological categories such as tissue domains[23] rather than predicting the genome-wide transcriptional state of individual cells within a tissue across space and time.

The scaling of spatial genomics measurements provides a new data source for capturing the structure of cell–cell interactions within a tissue and constructing predictive models[24]. Increasingly, spatial transcriptomic methods profile the molecular state of millions of single cells within the context of an intact human tissue[25, 26, 27, 28, 29, 30, 31]. Across diverse distributions of physiological states, spatial measurements resolve thousands of transcripts per cell, across targeted panels of hundreds to thousands of genes or genome-wide, implicitly recording the impact of cell–cell communication on the many human cell types present in an in vivo environment[13, 32, 33]. Methods for inferring cell–cell interactions from spatial data have focused on identifying which signals and pathways are engaged between adjacent cells[34, 35, 36, 37, 38]. Spatial genomics data motivates the development of computational methods that can convert observational measurements of cellular transcriptional states within human tissues into forward computational models of cell–cell signaling and tissue behavior.

Here we introduce the Cell Interaction Foundation Model (CIFM), a virtual tissue model that infers the language of cell interactions from spatial genomics data collected from primary human samples and enables auto-regressive forward simulations of tissue state across arbitrary, combinatorial gene expression perturbations. CIFM is enabled by three technical advances. First, CIFM is trained using masked transcriptome modeling, where individual cells are masked and a neural network predicts the transcriptional profile of a masked cell from the transcriptional state of neighboring cells in the tissue[39, 40, 41]. The strategy is analogous to masked language modeling but applied to cell state vectors and cell neighborhoods rather than word tokens and word sequences[39]. Second, at the architecture level, we develop a graph neural network where a gene expression encoder encodes the transcriptional state of cells in a neighborhood and a message passing layer learns to predict interactions between spatially proximate cells[42, 43]. Third, we develop a generative Monte-Carlo play-out strategy to propagate masked cell-state inference across a tissue in the background of therapeutic perturbations[44, 45]. Together, the advances enable CIFM to generate an explicit representation of cell state and cell–cell communication. Iterative resampling of the learned model plays the interactions out to simulate whole tissues under perturbation (Fig. 1).

**Figure 1.**
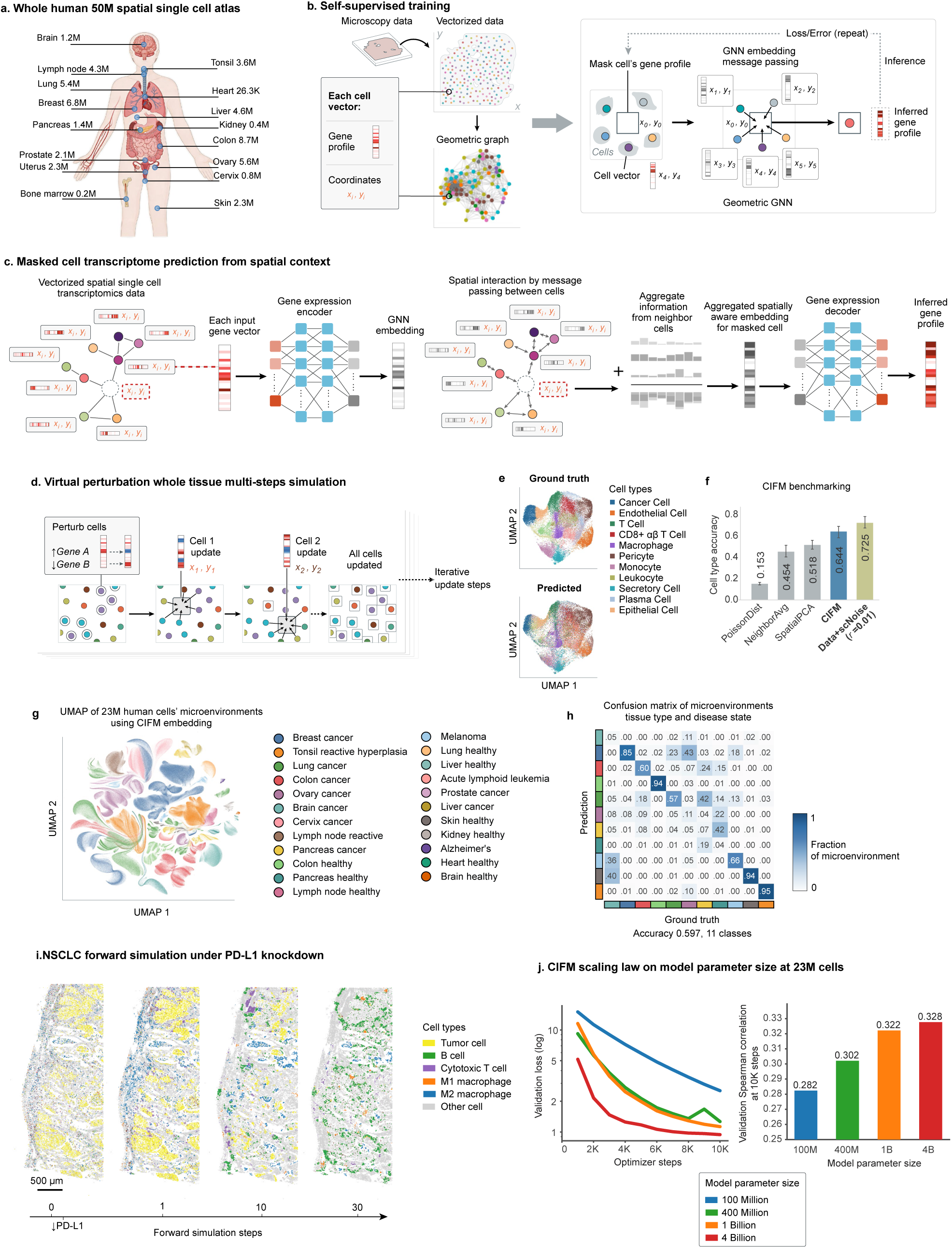
A geometric graph neural network trained by masked transcriptome prediction models cell interactions across human tissue. a, Curated spatial genomics corpus, showing cell counts by tissue. b, Self-supervised training. Spatial transcriptomics data are vectorized into per-cell records comprising a gene-expression profile and spatial coordinates and assembled into a geometric graph. The gene profile of a target cell is masked, a geometric graph neural network propagates information from neighboring cells by message passing, and the masked profile is inferred; reconstruction error is back-propagated. c, Masked cell transcriptome prediction. Each input gene vector is embedded by a gene-expression encoder; spatial interaction is modeled by message passing between neighboring cells; neighbor information is aggregated into a spatially aware embedding for the masked cell, from which a decoder reconstructs the genome-wide profile. d, Virtual perturbation and multi-step simulation. Selected genes are up- or down-regulated in selected cells, after which all cells are iteratively updated from their current neighborhoods, propagating the perturbation through the tissue. e, UMAP of measured (top) and CIFM-predicted (bottom) transcriptional profiles from held-out tissue, colored by annotated cell type. Cell clustering is performed based on expression profiles, and the resulting clusters are mapped and visualized in the UMAP space generated from ground-truth expression, with 54% of cells mapped to overlapping clusters. f, Cell-type annotation concordance for profiles predicted by CIFM and by three baselines — a Poisson expression model, neighbor averaging and SpatialPCA. Data+scNoise (r = 0.01) denotes measured expression perturbed by technical noise at 1% of signal and defines the attainable ceiling. g, UMAP of CIFM microenvironment embeddings for all the 23 million cells’ microenvironment (56 samples across Xenium V1, Xenium Prime and Visium HD), colored by the tissue and disease state of the source sample (23 categories). Each cell is represented by the 1,024-dimensional embedding of its neighborhood computed with the central cell masked. The vector describes the microenvironment of a cell rather than the cell itself. h, Tissue and disease state predicted from the same embeddings, evaluated by leave-one-sample-out cross-validation over the 44 samples belonging to the 11 classes represented on more than one sample. Columns are ground-truth classes and are normalized to one, so diagonal entries are per-class recall; axis colors follow the legend in g. Accuracy 0.597, against 0.091 expected by chance. i, Forward simulation of a non-small-cell lung carcinoma section under simulated PD-L1 knockdown, shown at simulation steps 0 (measured), 1, 10 and 30 and colored by cell type. Scale bar, 500 µm. j, Validation loss against optimizer steps for models of 100M, 400M, 1B and 4B parameters trained on a common pool of 23 million microenvironments. Validation Spearman correlation after 10,000 optimizer steps for the same four models on the same pool.

Trained on 23 million cell micro-environments, CIFM reproduces measured tissue behavior accurately enough to serve as a design tool, predicting held-out cell states near the noise ceiling of the measurements while also recapitulating perturbation responses measured in organoids, in vivo, and in engineered cell pairs. CIFM embeddings can be used for conventional spatial genomics tasks like patient classification, but the most important application of CIFM is its ability to forward simulate tissue behavior. Across cancer and auto-immune disease[46], we apply CIFM to screen over a million mono- and combinatorial perturbations, enabling global design of strategies that satisfy multiple therapeutic objectives simultaneously. Our computational screens identify a series of therapeutic concepts in cold tumors and point to an important role for Activin C in immune suppression that we validate in cell based assays[47]. CIFM also enables dynamic analysis, predicting transient and sustained responses in specific patients and supporting the design of dynamic therapeutic protocols to steer inflamed IBD tissue to a healthy state without overshooting. Broadly, CIFM shows that generative machine learning paradigms can enable the construction of human tissue foundation models, providing a new framework for forward tissue simulation and therapeutic design as well as a path to virtual organism level models.

## Results

### CIFM predicts single-cell transcriptional states using geometric graph neural networks

To build predictive models of tissue state and perturbation response, we developed the Cell Interaction Foundation Model (CIFM), an AI foundation model based on geometric self-supervised learning that can be trained on large spatial genomics datasets to model interactions across cells within tissues and to predict how cellular interactions modulate the distribution of gene expression states of cells within a tissue under normal and perturbed conditions. Conceptually, CIFM predicts the genome-wide transcriptional state of a cell within a tissue based on the states of its neighboring cells (Fig. 1b). The model is trained in a self-supervised masked transcriptome task in which individual cells within a spatial dataset are masked and the model is trained to predict the full transcriptional profile of each held-out cell. Following training and validation, the model can be used to predict the transcriptional state of cells following therapeutic perturbations to gene expression in single cells, and can therefore be used for therapy design (Fig. 1d).

Technically, CIFM represents local cell neighborhoods as a graph, with node features of gene expression and spatial coordinates, and edges representing spatial proximity between cells. To generate predictions of high-dimensional genome-wide transcriptional states from cell neighborhood graphs, we implemented an equivariant geometric graph neural network (GNN) architecture (GeoGNN) (Fig. 1c). The GNN architecture allows us to generate high-dimensional predictions on geometric graphs in which nodes carry vectorized single-cell expression data and edges represent spatial relationships between cells. The GeoGNN comprises three components. A gene expression encoder first embeds the genome-wide transcriptional state of each cell into a latent representation, so that the high-dimensional expression profile of every neighboring cell is compressed into a form on which the graph can operate. Message-passing layers then model communication between spatially proximate cells, propagating information along the edges of the neighborhood graph so that the representation of each cell is updated by the states of the cells around it; the information arriving at the masked position is aggregated into a single spatially aware embedding. A gene expression decoder finally maps that embedding back to a genome-wide profile, yielding the predicted transcriptional state of the masked cell.

We emphasize that CIFM predicts the global transcriptional state of each cell within a tissue based upon the global transcriptional state of neighboring cells. The architecture is equivariant, meaning that predictions are not altered by natural symmetries within the tissue, including spatial rotation and translation of the field of view. During training, the output of the GeoGNN is used to compute a mismatch loss as the combination of mean squared error (MSE) and binary cross entropy (BCE) against ground-truth transcriptional states, averaged across cell neighborhoods within samples and across different samples. We trained models with 100M, 400M, 1B and 4B parameters on GPU and TPU hardware with batch sizes adjusted for parameter count, each for a number of epochs determined through analysis of training and validation loss curves.

We trained CIFM on curated spatial genomics data and also developed an agentic system that continuously surveys the literature to identify and curate new datasets so that the CIFM data set is growing continuously in time. For our current release the model has been trained on data comprising 23 million cell micro-environments with transcriptional and spatial profiles from healthy and diseased samples across 4 platforms and 16 tissue types, with transcriptional states covering 18,000 measured genes. Each microenvironment is a single anchor cell together with the cells spatially adjacent to it, so that the corpus contains one microenvironment per profiled cell and every cell serves in turn as the masked target and as context for its neighbors. The model can be extended to other data sources including proteomic data. We trained CIFM on the full corpus to converged validation loss (Fig. 1j), indicating that it has learned consistent and stable cell interaction–state patterns from data at scale and is able to generate predictions on new cell neighborhoods, enabling downstream applications such as therapeutic target design.

The CIFM training task naturally extends to therapeutic prediction. We can predict the impact of a therapeutic perturbation on a tissue by manipulating gene expression vectors within the tissue and then performing masked transcriptome prediction on nearby cells. During training the model learns to infer the transcriptional state of a masked cell from the states of the cells surrounding it, and because every cell in the corpus serves in turn as the masked target, it learns this mapping across a range of cell neighborhood configurations. To predict the effect of an intervention, we edit the expression vectors of cells in a neighborhood to represent the intervention. Then, the target cell is masked as in training, and the forward pass returns the cell’s predicted transcriptional state under the new context.

### Cell neighborhoods predict single-cell transcriptional state

An open question in biology is how strongly a cell’s neighborhood and local interactions within a tissue influences its transcriptional state. Neighborhood effects have been characterized in specific contexts such as creating pro and anti-inflammatory signaling environments in cancer and autoimmune disease, but not systematically. CIFM allows the question to be asked quantitatively, by measuring how accurately the global transcriptional state of a cell can be predicted from the states of its neighbors alone (Fig. S1a).

We find that neighborhood context alone reconstructs the global structure of tissue expression. In a lung cancer sample, predicted and measured cell-by-gene matrices carry similar gene-program organization, with 54% of genes assigned to overlapping programs by Leiden clustering, against 56% for measured expression perturbed by noise (Fig. S1b); clustering cells on their predicted profiles recovers the measured cell clusters, with 54% of cells assigned to overlapping clusters against 64% for measured expression perturbed by noise (Fig. 1e). Predictions are also spatially faithful. Classifying randomly sampled regions as high or low for CD8A, MKI67 and T cell abundance, CIFM reaches F1 scores of 0.64, 0.75 and 0.80, in each case exceeding SpatialPCA[48] at 0.53, 0.62 and 0.72, whose predictions visibly smooth spatial structure rather than reconstructing it (Fig. S1c). Confusion matrices between predicted and measured cell clusters show correspondingly tighter diagonal structure for CIFM than for SpatialPCA (Fig. S1c).

Across samples, gene expression prediction accuracy approaches the limit set by underlying measurement noise. Averaged over eight tissues spanning skin, cervical cancer, prostate, ovarian cancer, lung cancer, ovary, reactive lymph node and breast cancer, mean squared error against ground truth was 0.689 for CIFM, against 0.773 for SpatialPCA, 0.875 for neighbor averaging where an error of 0.517 is obtained by measured expression perturbed by simulated single-cell sequencing noise at 10% of signal (Fig. S1d). Mapping predicted and measured profiles to cell types with a published annotation network, scTab[49], across 180 fine-grained categories and scoring two annotations as concordant when their most confident predictions intersect, 64% of cells were annotated identically from neighborhood-determined and ground-truth expression, against 52% for SpatialPCA, 45% for neighbor averaging and 15% for the Poisson model, and 73% for measured expression perturbed by noise at 1% of signal (Fig. 1f). CIFM therefore recovers 89% of the cell-type accuracy available in the data, and the residual gap is dominated by assay stochasticity rather than by model error.

### CIFM embeddings organize cell microenvironments by tissue and disease state

CIFM can also be applied to perform conventional tasks like tissue microenvironment origin or disease state classification. The forward pass of CIFM that predicts a masked transcriptome also produces an explicit vector representation of the neighborhood that generated the prediction, and that representation can be used for the classification tasks that spatial genomics conventionally addresses. We embedded the entire 23-million-cell atlas, masking each central cell so that the vector describes a cell’s surroundings rather than the cell itself, and projected the resulting representations into UMAP space (Fig. 1g). Microenvironments organize by tissue of origin and by disease state without supervision, and healthy and malignant tissue from the same organ occupy distinct regions of the embedding. Trained to predict tissue and disease state from these representations and evaluated by holding out entire samples, a classifier assigns held-out microenvironments to the correct one of eleven classes with an accuracy of 0.597, against 0.091 expected by chance (Fig. 1h).

### Prediction accuracy scales with model size

We next asked how prediction accuracy depends on the parameter size and information capacity of the model. We trained models of 100M to 4B parameters on a common pool of 23 million microenvironments (Fig. 1j) and measured performance. Larger models converged faster and with lower error, and the ordering was preserved throughout training (Fig. 1j). Similarly, validation correlation at 10,000 steps rose from 0.282 at 100M parameters to 0.302 at 400M, 0.322 at 1B and 0.328 at 4B (Fig. 1j). Accuracy therefore improves steeply up to a few hundred million parameters and then flattens, and we selected a 1B-parameter model as the operating point beyond which additional capacity returned little. This saturation is consistent with the task being limited by the data rather than by the model. The benchmark above places CIFM at 89% of the cell-type accuracy attainable from measured expression perturbed by technical noise (Fig. 1f), so the residual error that additional capacity would have to explain is largely assay stochasticity rather than unlearned biological structure.

### Autoregressive simulation propagates perturbations across a whole tissue

Therapeutic responses unfold over time, raising a central design question regarding how can interventions be chosen to produce sustained responses in chronic disease, as opposed to the transient responses appropriate to acute conditions. We extended CIFM to predict tissue-state trajectories by autoregressive simulation, feeding the output at step k as the input to step k+1. At each simulation step all cells are updated through sampling where each is masked and its genome-wide transcriptional state predicted from its current neighborhood, so that every cell’s update propagates to its neighbors at the following step (Fig. 1d, S5a). We often run simulations for 30 steps, which is sufficient to reach stable cell-state distributions in our samples. Simulation steps are ordinal and are not calibrated to physical time; all dynamic claims below concern the ordering of events rather than their explicit temporal rate.

Applied to a non-small-cell lung carcinoma section under simulated PD-L1 knockdown, iterating this update visibly restructures the tissue. M2 macrophages expand within the first step, cytotoxic T cells appear as discrete foci by step 10, and the tumor compartment is progressively replaced by infiltrating B and T cells by step 30 (Fig. 1i). The tissue is therefore reorganized by the propagation of a single perturbation through cell–cell interactions.

Applied to prostate cancer under simulated NSD2 knockdown, the simulation evolves the tissue at single-cell, whole-genome resolution, with cell-type composition shifting from a tumor-dominated state through transient myeloid and B cell responses (Fig. 2b,d). Gene programs separate into distinct temporal classes, including transient activation, sustained repression and low response, with AR/luminal programs rising early and decaying, interferon and antigen-presentation programs rising later and persisting, and neuroendocrine and cell-cycle programs progressively suppressed (Fig. 2c,d). The ordering of these programs was not supervised and is not present in any single training example, indicating that the model has learned dependencies between cell populations rather than static correlations.

**Figure 2.**
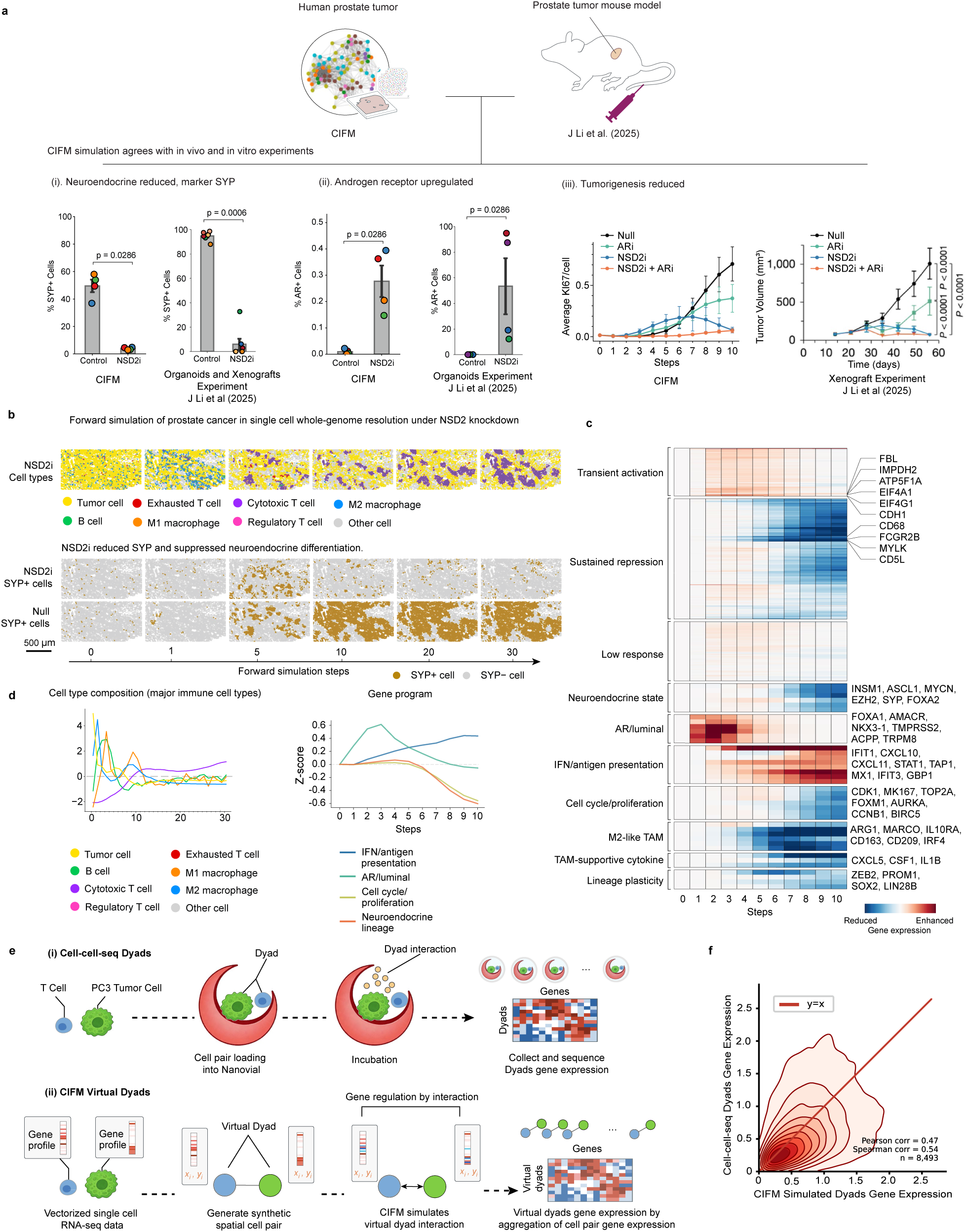
Simulated perturbation responses agree with responses measured in organoids, in vivo and in isolated cell pairs. a. CIFM simulation compared with published experiments in prostate cancer. (i.) Neuroendocrine differentiation, measured as the percentage of SYP+ cells, following NSD2 inhibition in simulation (left; p = 0.002) and in patient-derived prostate cancer organoids (right; p = 0.015). (ii.) Androgen receptor activity, measured as the percentage of AR+ cells, in simulation (left) and in the same organoids (right); AR+ cells are more abundant under NSD2 inhibition than under control in both. (iii.) Response to AR inhibition (ARi), NSD2 inhibition (NSD2i) and their combination, as simulated proliferation (average Ki67 per cell, left) and measured tumor volume in vivo (right). P values, Mann-Whitney U-test. b. Forward simulation of prostate cancer under simulated NSD2 knockdown at single-cell, whole-genome resolution, shown at simulation steps 0 (measured), 1, 5, 10, 20 and 30. Top, cells colored by type. Bottom, the same fields colored by SYP status under NSD2 knockdown (NSD2i) and under the unperturbed control simulation of the same tissue (Null). The two arms are comparable to step 5, after which the SYP+ compartment expands across the tissue in the control and remains sparse under NSD2i. Scale bar, 500 µm. c. Gene-program response over the first 10 simulation steps, grouped by temporal class (transient activation, sustained repression, low response) and by program identity; representative genes at right. d. Left, z-scored cell-type composition trajectory over 30 simulation steps. Right, z-scored activation trajectories for four gene programs over 10 steps. e. Experimental and virtual cell dyads. (i.) Cell–cell sequencing: single T cell–PC3 tumor cell pairs are co-loaded into nano-vials, incubated and sequenced. (ii.) CIFM virtual dyads: vectorized single-cell RNA-seq profiles are placed in synthetic spatial adjacency, their interaction simulated, and cell-pair expression aggregated. f. Gene expression in measured versus simulated dyads (n = 8,493; Pearson r = 0.47, Spearman *ρ* = 0.54). Red line, y = x.

### Simulated perturbation responses match measured responses in prostate cancer

We next tested whether simulated responses agree with experiment[50]. In prostate cancer, simulated NSD2 inhibition suppressed the neuroendocrine compartment, reducing the fraction of SYP-positive cells (p = 0.002), matching the suppression measured in patient-derived prostate cancer organoids (p = 0.015) (Fig. 2a(i)). The suppression is spatially resolved and runs against the unperturbed control. In the control simulation of the same tissue the SYP-positive compartment expands to cover much of the section from step 10 and persists to step 30, whereas under NSD2 inhibition it remains sparse throughout (Fig. 2b, bottom). The simulation also recovered the reciprocal trend in androgen receptor activity, with AR-positive cells more abundant under NSD2 inhibition than under control, as in the organoids (Fig. 2a(ii)). We then simulated four treatment arms: untreated, AR inhibition, NSD2 inhibition and the combination. We compared predicted proliferation with measured tumor growth in vivo. CIFM recovered both the ordering and the separation of the four arms, with the combination most strongly suppressive and AR inhibition intermediate (Fig. 2a(iii)).

### Learned interactions recapitulate physically measured cell–cell interactions

The preceding results test tissue-level outcomes; we next tested the cell-cell interaction model directly. Using public cell–cell sequencing data[51], in which single T cell–tumor cell pairs are co-encapsulated in nanovials, incubated and sequenced as “dyads”, we obtained expression profiles attributable to a defined pairwise interaction (Fig. 2e(i)). We generated matched virtual dyads by placing single-cell profiles of the same cell types in synthetic spatial adjacency and simulating their interaction with CIFM (Fig. 2e(ii)). Across 1,000 dyads on 8,493 genes, simulated and measured expression were correlated (Pearson r = 0.47, Spearman *ρ* = 0.54), with the relationship tracking the x=y line (Fig. 2f).

Consistent with this, CIFM-predicted perturbation responses cluster by receptor specificity. Aggregating single-gene over-expression predictions across seven Visium-HD tumor samples and mapping each target to its T cell binding partners in StringDB[52], targets binding the same T cell receptor produce similar genome-wide response patterns. For example, CXCR3 strong binders show significantly higher Pearson correlations among themselves than between binders and non-binders, and the same clustering extends to the targets of many other T cell molecules (Fig. S3a, S3b). These predictions are distinct from background data correlation, which relates perturbed-gene expression to CD8A expression only weakly (Pearson r = 0.14).

### Simulated cell injection produces responses corresponding to the identity of the added cells

The same machinery predicts the consequence of adding a cell rather than editing a gene. Injecting one hundred virtual cells carrying a cytotoxic CD8+ T cell marker profile into a lung cancer section suppressed MKI67 in the surrounding tissue over twenty simulation steps, whereas the same number of cells carrying a regulatory T cell profile, placed at identical positions, did not (Fig. S4), indicating that the predicted effect follows the transcriptional identity of the added cells rather than their presence.

### CIFM predicts genome-wide responses to perturbation of cell neighborhoods

Once trained, CIFM predicts how cells within a tissue respond to gene expression perturbations that alter cell–cell communication, enabling in silico therapeutic design at tissue scale. We performed virtual over-expression by setting the expression of target genes in cell neighborhoods to the maximum value in log1p-normalized space, and virtual knockdown correspondingly, then generated genome-wide predictions for cells in the perturbed neighborhoods (Fig. S2a). Predictions respect known signaling specificity and biochemical signal transduction. For example, the CIFM model predicts that over-expression of CXCL9, a chemokine that recruits CD8+ T cells through CXCR3, raises CD8A expression across a breast tumor, whereas over-expression of CXCL3, which does not bind CXCR3, does not (Fig. S2b).

Applying this across tumor samples to over 200,000 single and combinatorial perturbations of 487 genes, principally encoding intercellular signaling proteins, reveals that targets which up-regulate CD8+ T cells fall into cancer-type-general and cancer-type-specific groups (Fig. S2c). Perturbations cluster by the response they produce, and one cluster containing CXCL9, CXCL10, CXCL11, CCL18, CCL27, IL15, IL33 and PD-L1 (CD274) among others enhances CD8A response in all tumor types examined, with average effects ranging from 0.4 in melanoma to 1.5 in leukemia. Other perturbation clusters are strongly context-dependent. In cervical cancer most perturbation clusters reduce rather than enhance CD8A response, whereas the same perturbation clusters are neutral to positive in glioblastoma and ovarian cancer. Tumors therefore differ not only in the magnitude of their response but in its sign, and a target ranked highly in one cancer type cannot be assumed to transfer.

### Multi-objective design separates CD8+ T cell recruitment from regulatory T cell induction

Therapies must commonly satisfy several objectives at once, and genome-wide response prediction allows multi-objective design at scale. In cancer immunotherapy a central goal is to increase the abundance and activity of CD8+ killer T cells without also inducing immunosuppressive regulatory T cells. We therefore scored every perturbation on two axes simultaneously: enhancement of CD8A, marking CD8+ cytotoxic T cells, and reduction of FOXP3, marking immunosuppressive regulatory T cells. Using the CIFM model, we searched for perturbations occupying the quadrant that satisfies both (Fig. S2d). Over-expression of CXCL9 up-regulated CD8A but also up-regulated FOXP3, indicating a potential dual role in tumor tissue, whereas over-expression of CXCL10 or CXCL11 up-regulated CD8A more selectively. Combinatorial perturbations outperformed single-gene perturbations across the objective, so that manipulating two genes simultaneously produced both stronger CD8A enhancement and better separation from FOXP3, with CXCL10 combined with INHBC knockdown among the strongest, alongside CXCL10 with IGFBP7, EPO with TNFSF11 knockdown, and IL2 with IGFBP7 knockdown (Fig. S2d).

### Transient and sustained responses are distinguished by trajectory prediction

The CIFM model can be applied to analyze the dynamic trajectory of a therapeutic response and to design therapeutic strategies that generate transient or sustained responses for specific disease contexts. For example, in a breast tumor sample, CXCL9 over-expression raised CD8A expression by step 5 but the effect decayed by step 30, whereas PD-1 (PDCD1) knockdown alone produced little change; combining the two sustained the elevation through step 30 (Fig. S5b). Clustering all perturbations by their response trajectory resolves three groups that we name sustained, transient and minimal up-regulation. We find that sustained responses are dominated by combinations pairing a chemokine with knockdown of an inhibitory or angiogenic molecule, including CXCL9 or CXCL10 with PD-1 and VEGFA, CXCL11 with PD-L1 and VEGFA, and WNT5B with TNFSF11 (Fig. S5c). Transient up-regulation, by contrast, characterizes single-agent chemokine perturbations such as CXCL13 and IFNG. Ranking perturbations by their effect at step 30 rather than at their peak therefore reorders the candidate list substantially, and does so differently in different tumors. As an example, in luminal B breast cancer the highest-ranked strategies were ↑CXCL10,↓INHBC; ↑CXCL10,↓PD-1,↓VEGFA; ↓SFRP5; ↑CXCL10,↓PD-1; and ↑CXCL11,↓CYTL1, whereas in non-small-cell lung cancer they were ↑CXCL11,↓CYTL1; ↑WNT5B,↓TNFSF11; ↑CXCL9,↓PD-L1,↓VEGFA; ↓CSHL1; and ↓IGFBP6 (Fig. S5d).

### Simulated tissues execute an ordered immune recruitment cascade

We next asked whether simulation at whole-tissue scale reproduces the ordered multicellular dynamics that define an immune response, and whether the heterogeneity of those dynamics across patients recovers clinically recognized phenotypes. Simulating non-small-cell lung carcinoma under combined CXCL9 induction and PD-1 knockdown, the tissue passes through an ordered recruitment cascade over thirty steps where a type I and II interferon response with chemokine secretion rises first and peaks by step 4, myeloid activation and pro-inflammatory cytokine signaling follow and peak near step 8, and CD8+ T cell cytotoxicity and cytokine secretion rise late, by step 25, and persist to step 30 as cytotoxic T cells attract and then infiltrate (Fig. 3a). Cell-type composition tracks the same sequence, with tumor cells declining from step 5 onward as cytotoxic and regulatory T cells and B cells expand. Neither the ordering of these programs nor the delay between them was supervised, and no single training example contains a time course; the sequence arises from iterated application of a model trained only to infer a cell state from its neighbors.

**Figure 3.**
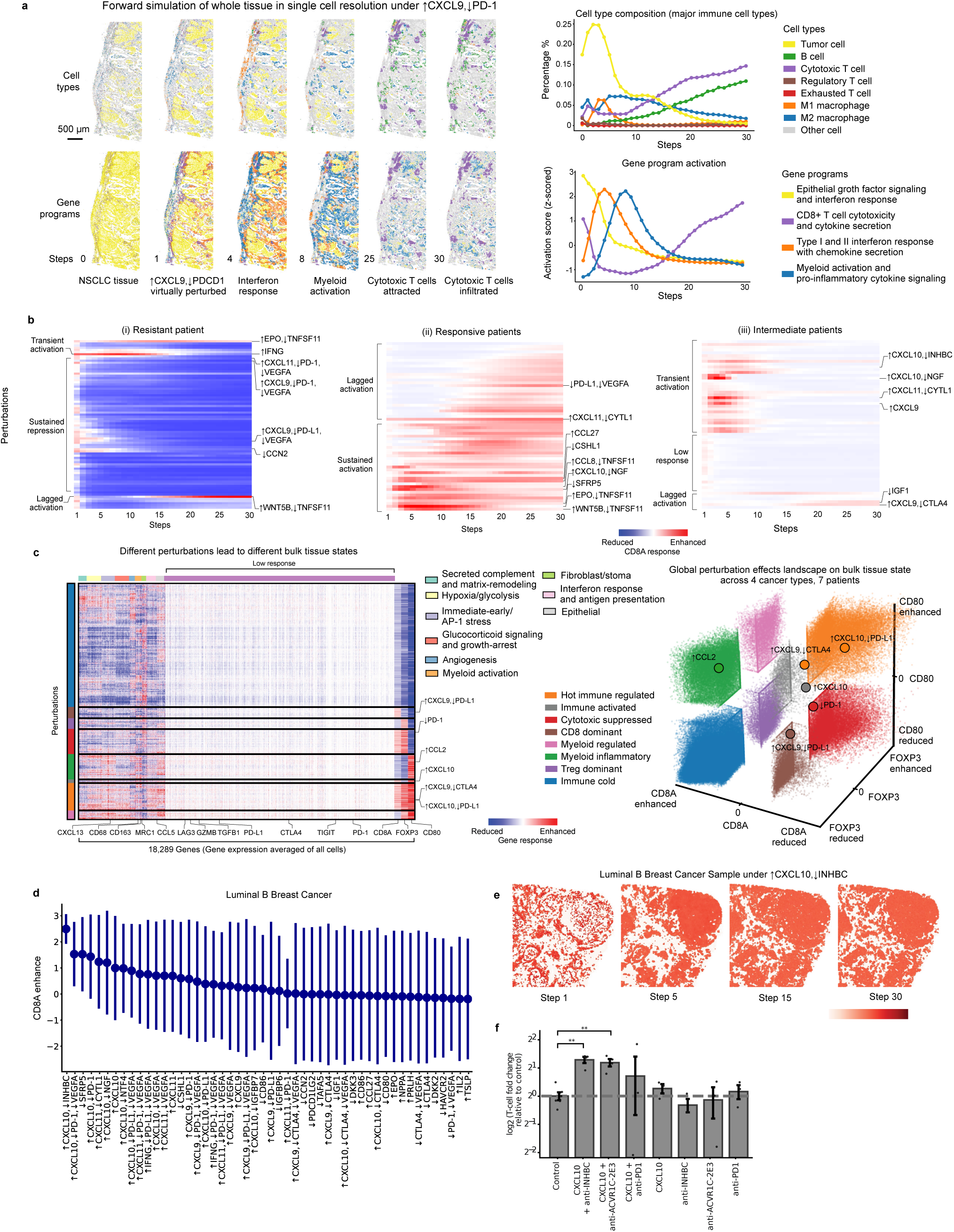
Forward simulation resolves patient-specific response classes, builds a tissue programming landscape and identifies a combination that expands T cells in vitro. a, Forward simulation of NSCLC tissue under combined CXCL9 induction and PD-1 knockdown over 30 steps, colored by cell type (top) and dominant gene program (bottom). Right, cell-type composition and z-scored gene-program activation, showing an ordered cascade of interferon response by step 4, myeloid activation by step 8, attraction of cytotoxic T cells by step 25 and their infiltration by step 30, accompanied by progressive loss of tumor cells. b, CD8A response trajectories across 30 simulation steps in (i) resistant, (ii) responsive and (iii) intermediate patients, grouped by temporal class; selected perturbations labeled. c, Left, genome-wide response across 18,289 genes averaged over all cells for each perturbation, annotated by tissue program; selected marker genes below. Right, every simulated tissue state across four cancer types and seven patients, embedded along axes of CD8A, FOXP3 and CD80 response and colored by destination state. d, Perturbations in luminal B breast cancer ranked by predicted CD8A enhancement at simulation step 30; points, mean; bars, s.d. across patients. e, Spatial CD8A response in a luminal B breast cancer sample at simulation steps 1, 5, 15 and 30 under the top-ranked perturbation, CXCL10 induction with INHBC knockdown. f, T cell expansion in co-culture as log2 fold change relative to control; points, replicates; bars, mean ± s.d.; *\* * P <* 0.01, t test.

### Patient tumors separate into resistant, responsive and intermediate classes

Applying the same procedure across patients reveals that identical interventions produce opposite tissue responses in different tumors. Ranking perturbations by their CD8A response trajectory separates patients into three classes (Fig. 3b). In resistant tumors the majority of perturbations drive transient or weak activation of CD8A, and even combinations that succeed at the cohort level, including CXCL9 or CXCL11 with PD-1 and VEGFA blockade, produce only transient activation before reverting in specific patients (Fig. 3b). In responsive tumors the same panel produces sustained activation, led by CXCL10 with NGF knockdown, SFRP5 knockdown and EPO with TNFSF11 knockdown, with PD-L1 and VEGFA blockade acting through a lagged trajectory that emerges only after step 10. Intermediate tumors show predominantly transient or low responses. The model therefore reproduces the division into responders and non-responders that limits checkpoint therapy in the clinic, and does so from tissue composition and architecture alone, without access to any treatment outcome.

### A tissue programming landscape relates interventions to destination states

Scaling the analysis produces a global map of how interventions move tissues between states. Across perturbations and 18,289 genes, responses partition into coherent tissue programs including interferon response and antigen presentation, myeloid activation, angiogenesis, hypoxia and glycolysis, secreted complement and matrix remodeling, glucocorticoid signaling and growth arrest, immediate-early AP-1 stress, fibroblast and stromal, and epithelial. Additionally, a large group of genes respond little (Fig. 3c). Embedding every simulated tissue state on axes of CD8A, FOXP3 and CD80 response, across four cancer types and seven patients, resolves eight destination states which we name immune cold, immune activated, hot immune regulated, CD8 dominant, cytotoxic suppressed, Treg dominant, myeloid regulated and myeloid inflammatory (Fig. 3c). Individual interventions occupy separable regions of this space, for example, CCL2 induction drives tissues toward myeloid states, CXCL10 activation with PD-L1 blockade toward CD8 dominance, CXCL9 with CTLA4 blockade toward a hot but regulatory-T-cell-infiltrated state, so that the landscape functions as a programming diagram relating an intervention to the tissue state it produces.

### A small set of interventions recurs across patients and tumor types

A small number of interventions recur at the top of this landscape across patients and tumor types. Ranking all perturbations in luminal B breast cancer by predicted CD8A enhancement places induction of CXCL10 with knockdown of INHBC first, ahead of CXCL10 combined with PD-1 and VEGFA blockade, CXCL11 with CYTL1 knockdown, and CXCL10 with NGF knockdown (Fig. 3d). Forward simulation of the top-ranked intervention shows the CD8A response beginning focally and spreading until it covers the tissue by step 30 (Fig. 3e).

### A designed CXCL10–INHBC combination expands T cells in co-culture

We used the trajectory screen to design an intervention in luminal B breast cancer, ranking all perturbations by predicted enhancement of CD8A at step 30 (Fig. 3d-f). The top-ranked perturbation was induction of CXCL10 combined with suppression of INHBC, an inhibin *β*-subunit signaling through the receptor ACVR1C, ahead of combinations involving PD-1 or CTLA4 blockade. Forward simulation showed the CD8A response emerging locally and spreading through the tissue over 30 steps (Fig. 3e).

We tested the top-ranked prediction experimentally. In co-culture, CXCL10 combined with an antibody against INHBC significantly expanded T cells relative to control, as did CXCL10 combined with an antibody against ACVR1C, the receptor through which the inhibin *β*C subunit signals (Fig. 3f). Neither CXCL10 alone, anti-PD-1 alone, nor CXCL10 combined with anti-PD-1 produced a significant increase in the same assay, nor did blockade of INHBC, ACVR1C or SFRP5 without CXCL10. In this way blocking antibodies against the ligand and against its receptor independently reproduce the effect, supporting the hypothesis that the CIFM prediction identified a suppressive signaling axis whose impact is conditional on the chemokine context the model specified. In the absence of chemokine the suppressive effect of INHBC is masked, but the combinatorial impact of chemokine activation plus INHBC inhibition is required to recruit immune cells into the tumor compartment while evading suppression by Activin signaling.

### CIFM extends to inflammatory bowel disease and predicts a coordinated program switch

To test whether a single model transfers across disease areas, we applied CIFM to colonic tissue from patients with ulcerative colitis and Crohn’s disease, fine-tuning the pre-trained model onto a 980-gene panel measured on a different spatial platform. The corpus comprises nine biopsies[53] — three healthy controls, three with ulcerative colitis and three with Crohn’s disease (Fig. 4a). We decomposed the measured tissue into nine transcriptional programs by orthogonal matrix factorization, spanning smooth muscle, fibroblast matrix, colonocyte barrier, epithelial repair, plasma cell immunoglobulin secretion, T cell activation, antigen presentation, myeloid cytokine and mast cell signaling, and acute phase signaling[54], and used that fixed basis to read out every simulation that follows.

**Figure 4.**
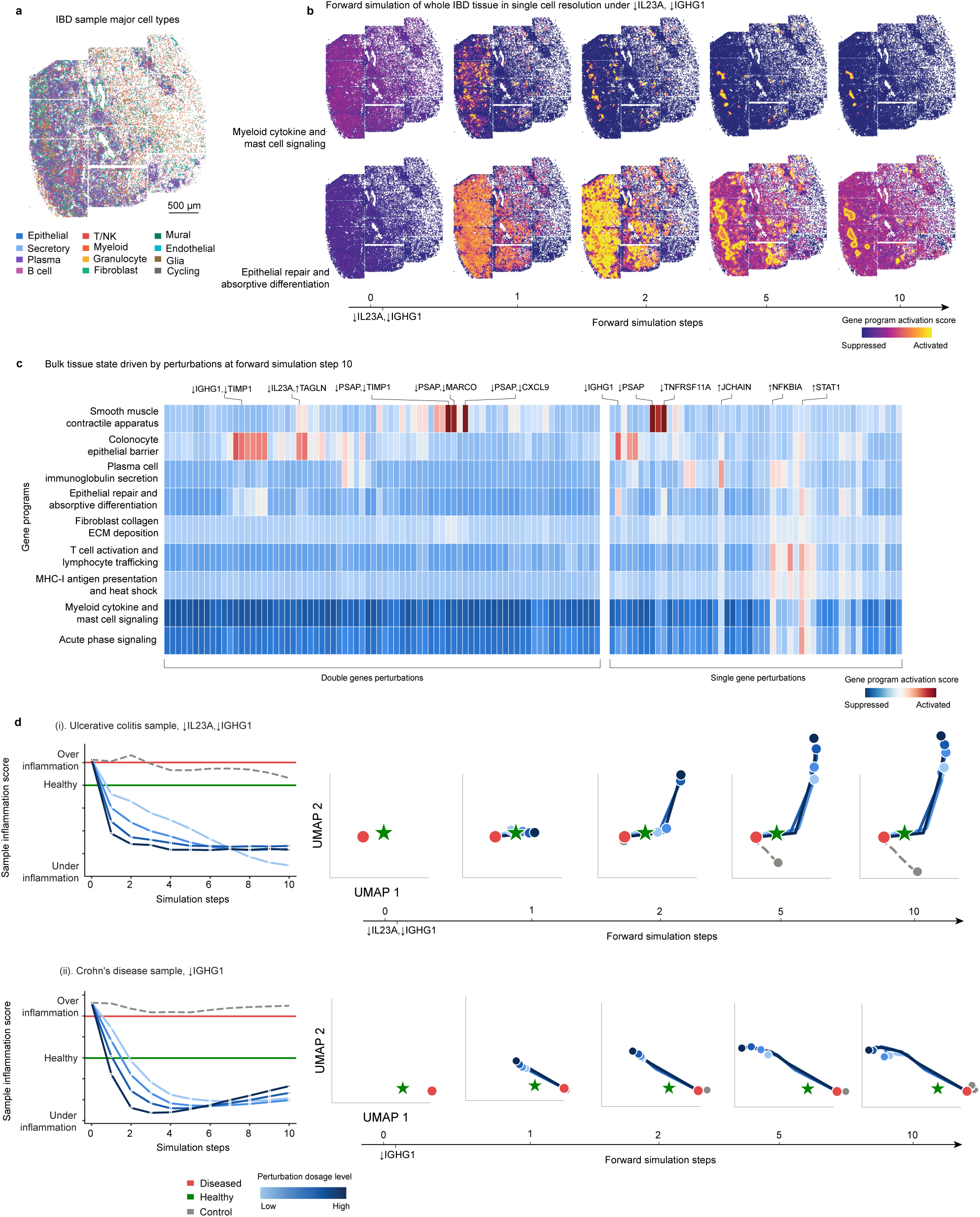
CIFM extends to inflammatory bowel disease and predicts overexposed perturbations driving inflamed colonic tissue past a healthy state. a, Measured cell types in an ulcerative colitis colon biopsy (980-gene panel; 35,086 cells), colored by twelve lineages collapsed from the published single-cell annotation. Scale bar, 500 µm. b, Forward simulation of the same tissue at single-cell resolution under combined IL23A and IGHG1 knockdown, at simulation steps 0 (measured), 1, 2, 5 and 10, colored by activation of the myeloid cytokine and mast cell signaling program (top) and the epithelial repair and absorptive differentiation program (bottom). Activation is the non-negative coefficient of a fixed nine-program orthogonal matrix factorization of the measured tissue; a single color scale is used throughout. c, Change in activation of nine tissue programs at simulation step 10 for 127 perturbations (76 two-gene, left; 51 single-gene, right), measured as the difference from the unperturbed simulation of the same tissue and averaged over three ulcerative colitis and three Crohn’s disease samples. Perturbations are ordered by hierarchical clustering within each block; selected perturbations labeled. ↓, knockdown; ↑, over-expression. d, Tissue inflammation score over ten simulation steps for (i) an ulcerative colitis sample under IL23A and IGHG1 knockdown and (ii) a Crohn’s disease sample under IGHG1 knockdown. The score is a logistic classifier separating healthy control from diseased tissue in the same nine-program space, computed on composition-normalized activations so that it is insensitive to overall transcriptional dilution, and rescaled so that 0 is the mean of the three healthy controls (green) and 1 the mean of the six diseased samples (red). Blue, four perturbation exposure levels; dashed gray, the unperturbed control simulation of the same tissue. Right, the same trajectories in a UMAP of program activations, fitted separately for each perturbation; red, untreated diseased tissue; green star, the mean of the three healthy controls; gray, the unperturbed control.

Simulated knockdown of IL23A together with IGHG1 produces a coordinated and spatially organized switch between two of these programs. Myeloid cytokine and mast cell signaling, one of the inflammatory of the nine programs, falls across the tissue within two simulation steps and continues to fall (Fig. 4b, top). Over the same interval epithelial repair and absorptive differentiation rises, most steeply between steps one and two, then flattens (Fig. 4b, bottom). Because both effects diverge from the baseline dynamics of the untreated simulation, we attribute the dynamics to the intervention rather than to the dynamics of iterated prediction.

Scoring 127 single- and two-gene perturbations against the unperturbed simulation of the same tissue, most of the program responses are suppressive, with the myeloid and acute phase programs suppressed most strongly (Fig. 4c). Ulcerative colitis and Crohn’s disease samples respond along the same axis but not with the same magnitude. The response profiles for the two diseases correlate at r = 0.90 while Crohn’s tissue responds 1.6-fold more strongly, so the model reproduces a shared response structure with disease-specific sensitivity. The screen prioritizes selective suppression of the IL-23 p19 subunit over TNF*α* or shared IL-12/23 p40 suppression in these tissues[55, 56]. Blockade of TNF*α* ranks 122nd of 127 by suppression of the myeloid program and blockade of the IL-12/23 p40 subunit 103rd, while the strongest interventions are combinations built on knockdown of the IL-23 p19 subunit, and adding TNF blockade to IL-23 blockade improves the predicted response. The model therefore predicts that TNF is a weak lever on myeloid inflammation in these tissues, consistent with the substantial fraction of patients who do not achieve durable remission on anti-TNF therapy[57]. We note that simulated suppression of a ligand transcript is not equivalent to antibody neutralization of the secreted protein, and that the ranking is therefore best read as a relative ordering of tissue-level consequence rather than as a prediction of clinical efficacy.

### Simulation identifies the exposure level and simulation steps that return inflamed tissue to a healthy state

Chronic inflammatory disease is usually treated over long periods[58], so the design problem is not only which target to engage but how strongly and for how long, since a perturbation that corrects a tissue can also carry it past the state it was meant to reach. Trajectory prediction turns that into a quantity CIFM can optimize. We projected each simulated tissue onto the single direction that best separates healthy control from diseased tissue in the nine-program space, calibrated so that zero is the healthy mean and one the diseased mean, and repeated every simulation at four perturbation exposure levels, giving for each intervention a surface over exposure level and exposure simulation steps (Fig. 4d). That surface is not monotone in either variable, so an optimum exists. In a Crohn’s disease sample under IGHG1 knockdown, an intermediate exposure level brings the tissue from 1.32 to 0.04 — the healthy value — after a single step of exposure, and then past it to *−*1.20 by step five; the weakest level reaches the same point after two steps, and the strongest overshoots within one (Fig. 4d(ii)). The tissue can therefore be placed at the healthy value by more than one combination of level and simulation steps, and missing that combination in either direction leaves it either still inflamed or over-suppressed. Across the same grid the unperturbed simulation of the same tissue never crosses either reference, so the trajectory is driven by the intervention and not by the dynamics of iterated prediction.

Searching both variables together is what makes the prediction useful, because the admissible region is narrow and depends on the tissue. The ulcerative colitis sample sets the opposite boundary where combined IL23A and IGHG1 knockdown crosses both references within a single step at every level tested and settles far past the healthy value, so the model predicts that this intervention has no admissible setting on the grid simulated and would have to be applied more weakly or more briefly still (Fig. 4d(i)). We are precise about what this axis measures. It is the direction along which healthy and diseased tissue are most separable, so reaching zero means reaching the healthy value of that projection rather than matching healthy tissue in every program, and the simulations hold the perturbation in place throughout, so they constrain where a given exposure lands the tissue rather than whether it remains there once exposure ends.

## Discussion

We developed CIFM, an AI foundation model that predicts how cell interactions determine cell state at single-cell, genome-wide resolution, and that supports computational design of therapeutic perturbation strategies at scale. The model rests on self-supervised masking and geometric graph neural networks, trained on 23 million cell microenvironments from diverse spatial genomics platforms. Applied to over 200,000 interventions targeting 487 signaling molecules, CIFM enables design against multiple objectives simultaneously. For example, the model confirms the simple prediction that CXCL9 increases T cell infiltration, as previously known[59], while revealing the more subtle and undesirable induction of regulatory T cells that accompanies it. The model identifies a combinatorial manipulation — CXCL10 with INHBC knockdown — that drives T cell recruitment without regulatory T cell induction. Extending the model to trajectory prediction distinguishes transient from sustained responses and shows that transient strategies can commonly be converted to sustained ones by a second perturbation. The top-ranked prediction of this analysis was validated in vitro, where it exceeded an anti-PD-1 combination.

## Author Contributions

Conceptualization, Y.L, Z.J.W, Y.Y, M.T; Methodology, Y.L, W.Z, J.G, Y.Y, M.T; Data Analysis, Y.L, J.Y, L.C, Y.Y, M.T; Experiments, Y.L, Y.C, J.Y, Y.P; Data Curation Y.L, J.Y, L.C, K.F, R.L, Y.Y; Writing, Y.L, Y.Y, M.T.; Visualization, Y.L, Y.Y, M.T; Interpretation: Y.L, Z.J.W, Y.P, Y.Y, M.T; Supervision, Y.Y, M.T; Funding Acquisition, M.T.

## Acknowledgements

This work was supported by Orr Family Foundation, the National Institutes of Health (TR01 GM150125, R01HD100039, R33CA297969 and R33CA247744 from the NCI, and the Information Technology for Cancer Research (ITCR) programme), the National Science Foundation Center for Cellular Construction (DBI-1548297), the Heritage Medical Research Institute, Charles Trimble, the Shurl and Kay Curci Foundation, the Merkin Institute for Translational Research, 10x Genomics, Amgen, and the Chan Zuckerberg Initiative.

---

**Figure S1.**
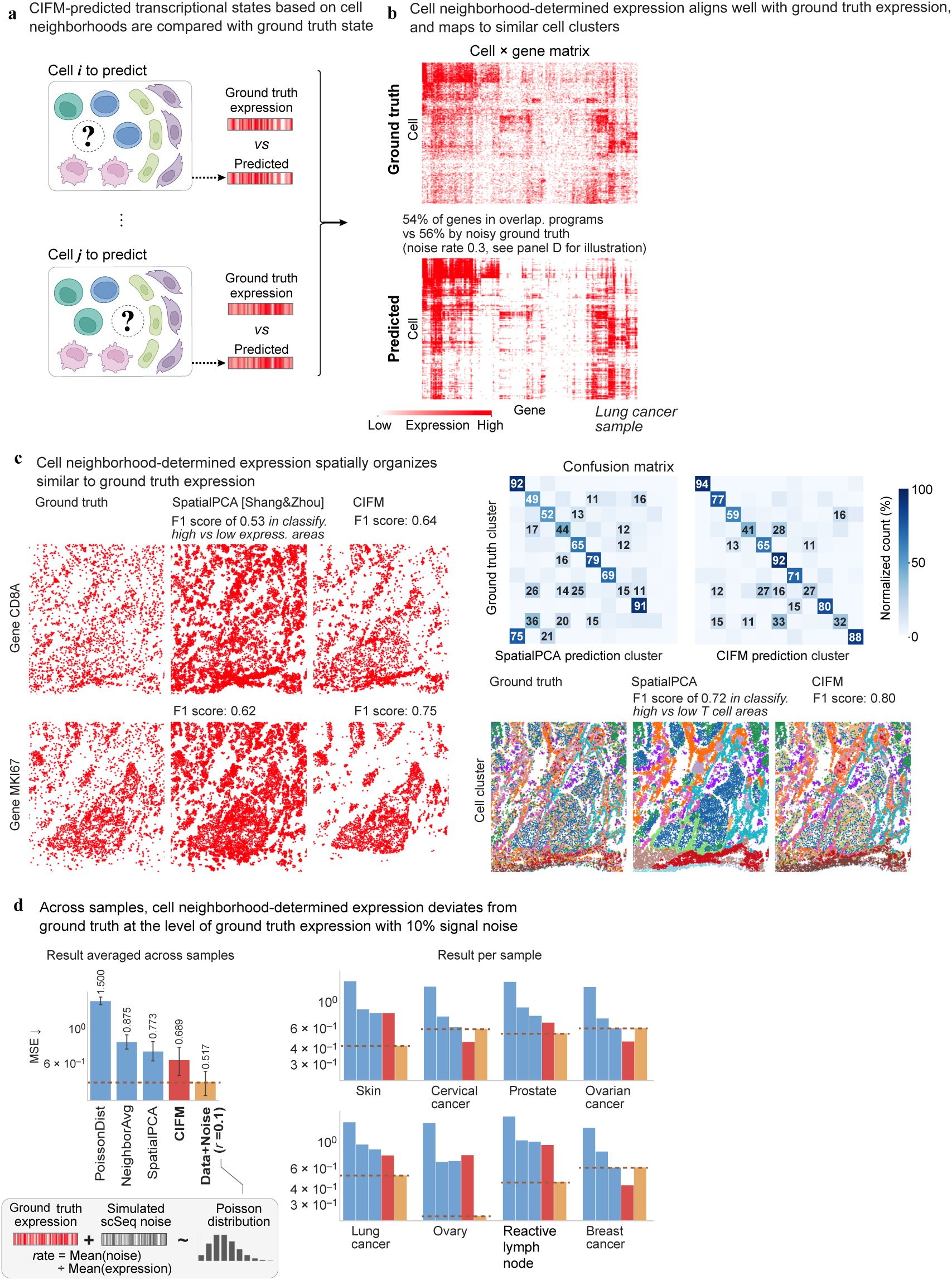
Cell neighborhoods determine single-cell transcriptional state. a, Evaluation design. For each held-out cell, the genome-wide transcriptional state predicted from the surrounding neighborhood is compared with the measured profile of that cell. b, Cell-by-gene matrices for measured (top) and predicted (bottom) expression in a lung cancer sample, with 54% of genes assigned to overlapping gene programs by Leiden clustering against 56% for measured expression perturbed by noise at a rate of 0.3. c, Spatial organization of measured, SpatialPCA-predicted and CIFM-predicted expression for CD8A and MKI67 and for cell-cluster identity. Classifying randomly sampled regions as high or low, CIFM reaches F1 scores of 0.64, 0.75 and 0.80 against 0.53, 0.62 and 0.72 for SpatialPCA. Right, confusion matrices between predicted and measured cell clusters for both methods, normalized counts. d, Mean squared error between predicted and measured expression averaged across samples (left) and per sample for eight tissues (right); dashed line, measured expression perturbed by simulated single-cell sequencing noise at 10% of signal. Inset, construction of the noise reference.

**Figure S2.**
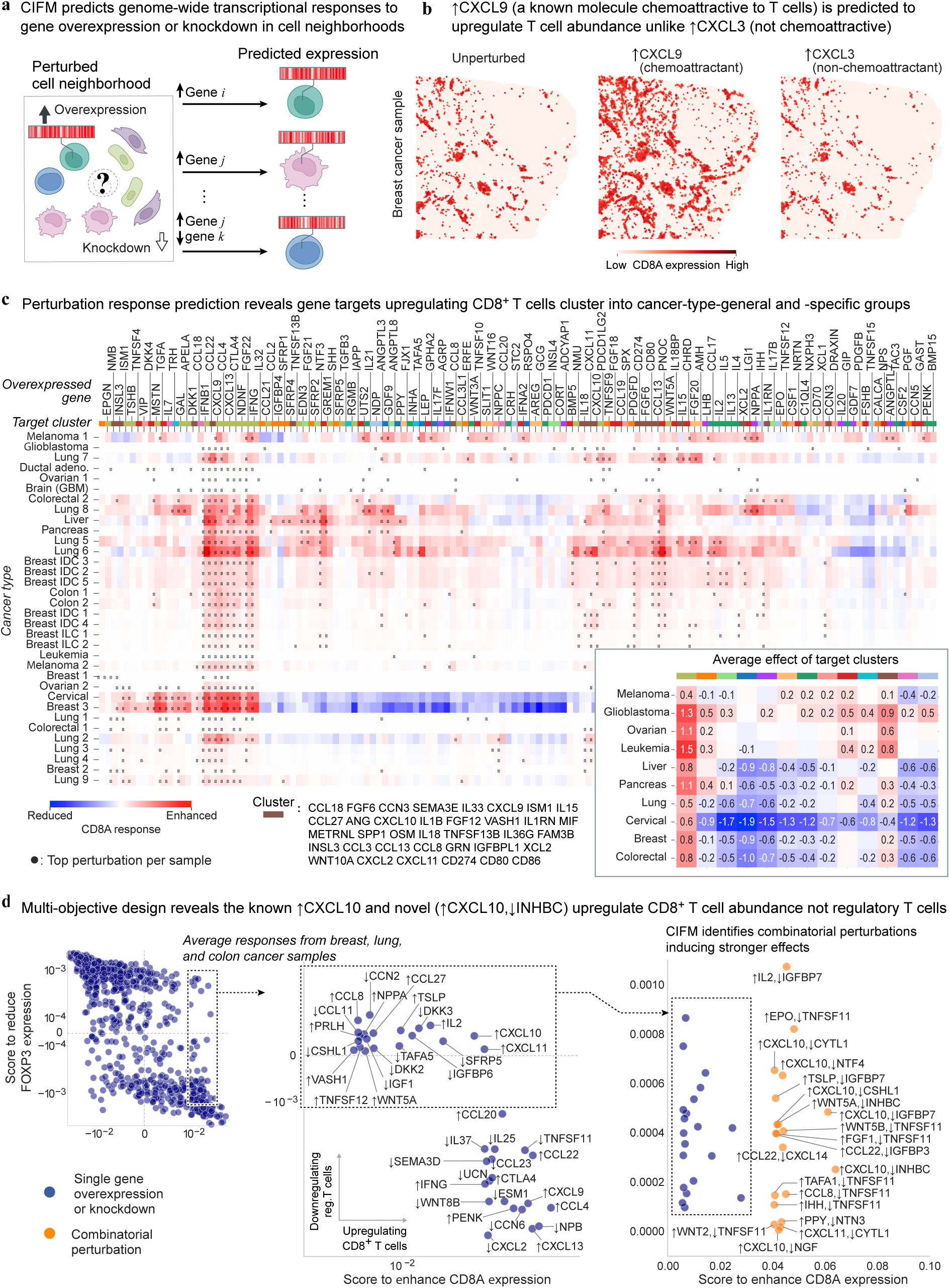
CIFM predicts genome-wide responses to perturbation and enables multi-objective design. a, Virtual perturbation. Genes are over-expressed or knocked down in the cells of a neighborhood and CIFM predicts the resulting genome-wide transcriptional state of the query cell, singly or in combination. b, Predicted CD8A expression across a breast cancer sample under no perturbation, CXCL9 over-expression and CXCL3 over-expression; CXCL9 binds CXCR3 and CXCL3 does not. c, Predicted CD8A response to single-gene over-expression across roughly thirty-four tumor samples spanning ten cancer types, with perturbations grouped into target clusters; dots mark the top perturbation per sample. Inset, average effect of each target cluster by cancer type. d, Every perturbation scored simultaneously for CD8A enhancement and FOXP3 reduction, with single-gene perturbations in blue and combinations in orange; successive panels magnify the quadrant satisfying both objectives.

**Figure S3.**
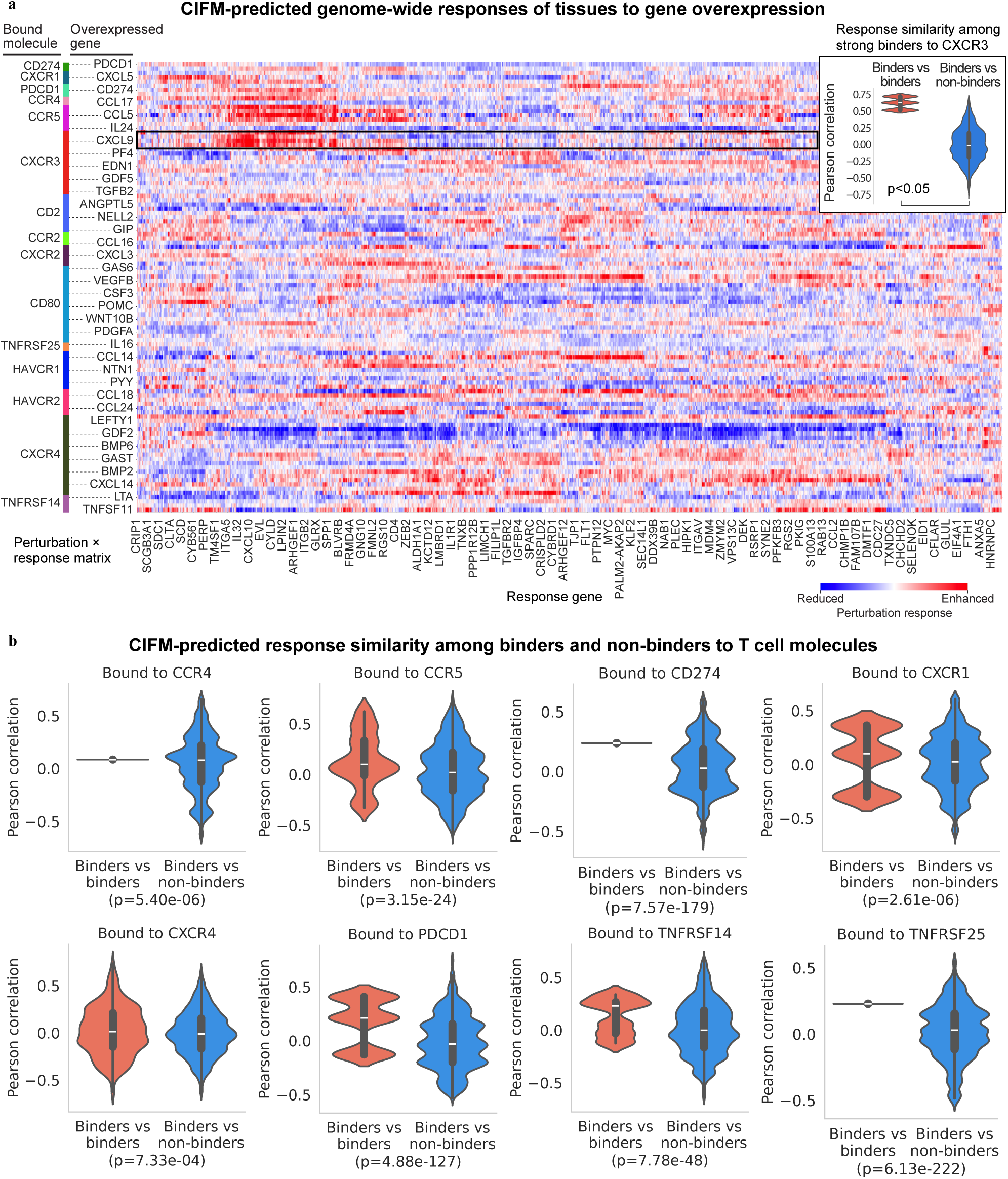
CIFM predicts that ligands binding the same T cell receptor produce similar genome-wide tissue responses. a, Genome-wide perturbation-response landscape. CIFM simulated saturating single-gene over-expression of 487 secreted-signaling and immune-checkpoint genes in cell neighborhoods from each of seven Visium HD human tumor sections (four lung, two breast, one colorectal). For each perturbation the response is the change, relative to the unperturbed simulation, in predicted expression of 18,289 genes; responses were z-scored across perturbations within each sample and averaged across samples. Rows, the 106 over-expressed ligands assignable to a T cell surface receptor in StringDB v12.0 (experimental-evidence score *≥* 0.05, each ligand assigned to its highest-scoring receptor, receptors with a single ligand excluded), grouped by bound molecule (left color bar, 15 receptors) and clustered within each group; columns, the 1,000 most responsive genes, hierarchically clustered. Color, response z-score clipped at *±*3. Every third row and every thirteenth column is labeled. Black box, the CXCL9 row. Inset, Pearson correlations between genome-wide response vectors for pairs of strong CXCR3 binders (CXCL9, CXCL10, CXCL11; experimental-evidence score *≥* 0.3; red) and for pairs formed between a CXCR3 ligand and a ligand of another receptor (blue). b, The same comparison for each receptor reaching significance (8 of 15 tested). Red, 1,000 randomly sampled pairs of ligands of the same receptor; blue, 14,000 pairs formed between a ligand of that receptor and a ligand of each of the other fourteen receptors; values are Pearson correlations across 18,039 named genes. P values, one-sided two-sample t test (binders *>* non-binders), d.f. = 14,998. Ligands per receptor: CCR4 2, CCR5 6, PD-L1 2, CXCR1 3, CXCR4 19, PD-1 3, TNFRSF14 4, TNFRSF25 2.

**Figure S4.**
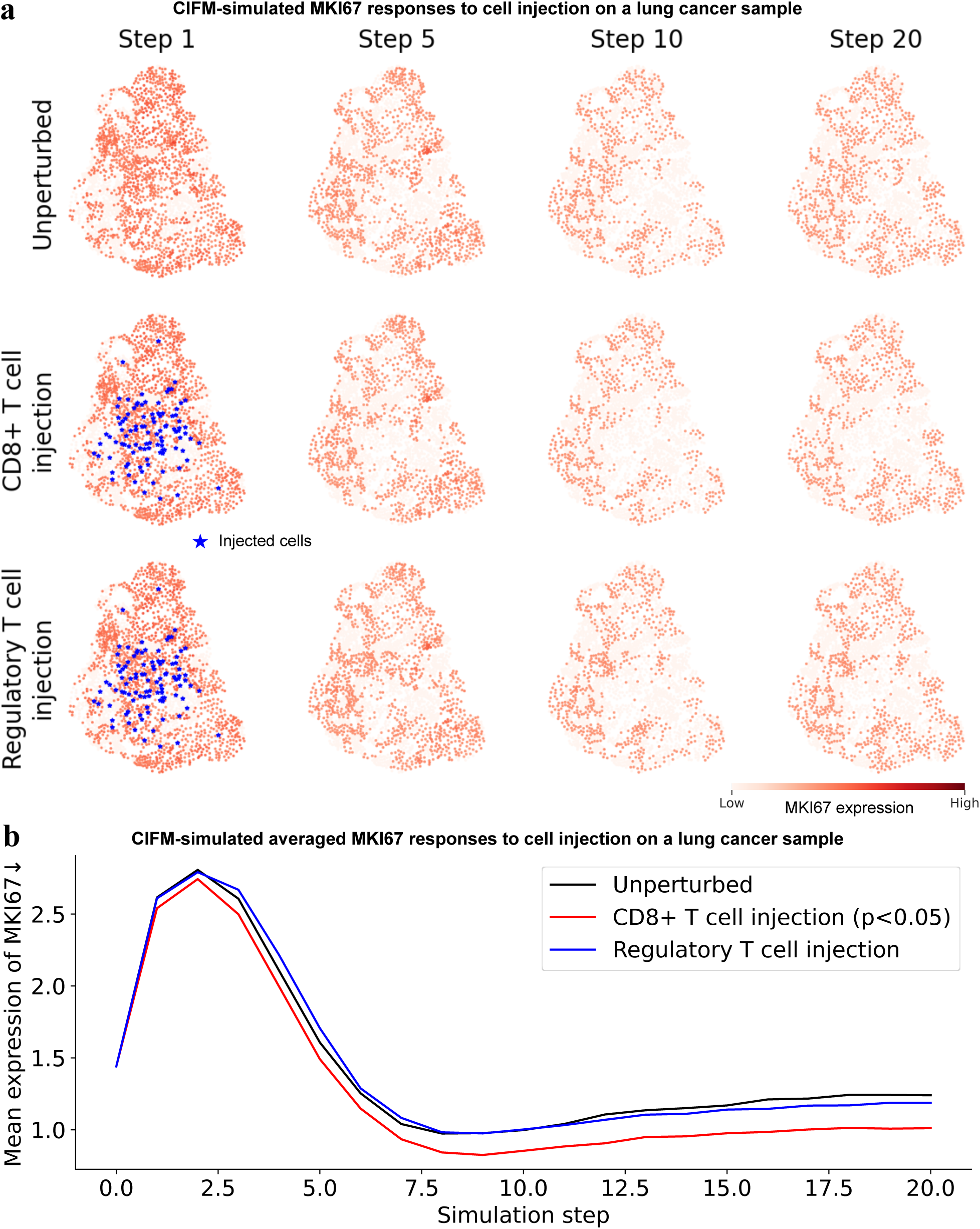
Virtual injection of cytotoxic but not regulatory T cells suppresses proliferation in simulated lung tumor tissue. a, CIFM simulation of virtual T cell injection into a Xenium V1 human lung cancer section (160,448 cells, 480-gene panel). One hundred virtual cells carrying a marker-defined synthetic transcriptome — effector-memory CD8+ T cell (CCL5, CD8A, NKG7, GZMK, CD3D, GZMA) or regulatory T cell (CTLA4, FOXP3, IL2RA, CD3D, CD247) — were placed at Gaussian-distributed positions (s.d. 100 µm) around the centroid of the section, and the tissue was evolved for 20 update steps; injected cells were held at their initial profile throughout. Panels show MKI67 expression in the 2,326 endogenous cells lying within 100 µm of an injected cell, at steps 1, 5, 10 and 20, for unperturbed tissue (top), CD8+ T cell injection (middle) and regulatory T cell injection (bottom). Blue stars, positions of injected cells. b, Mean MKI67 across the same 2,326 cells against simulation step. The transient rise and fall over the first ten steps occurs in the unperturbed arm as well and is a property of the iterative update; the arms separate from approximately step 7. CD8+ T cell injection reduces MKI67 relative to unperturbed tissue at the final step (one-sided paired Wilcoxon signed-rank across 2,326 cells, *P* = 4 *×* 10*^−^*^33^), whereas the same number of regulatory T cells at the same positions does not (*P* = 0.90).

**Figure S5.**
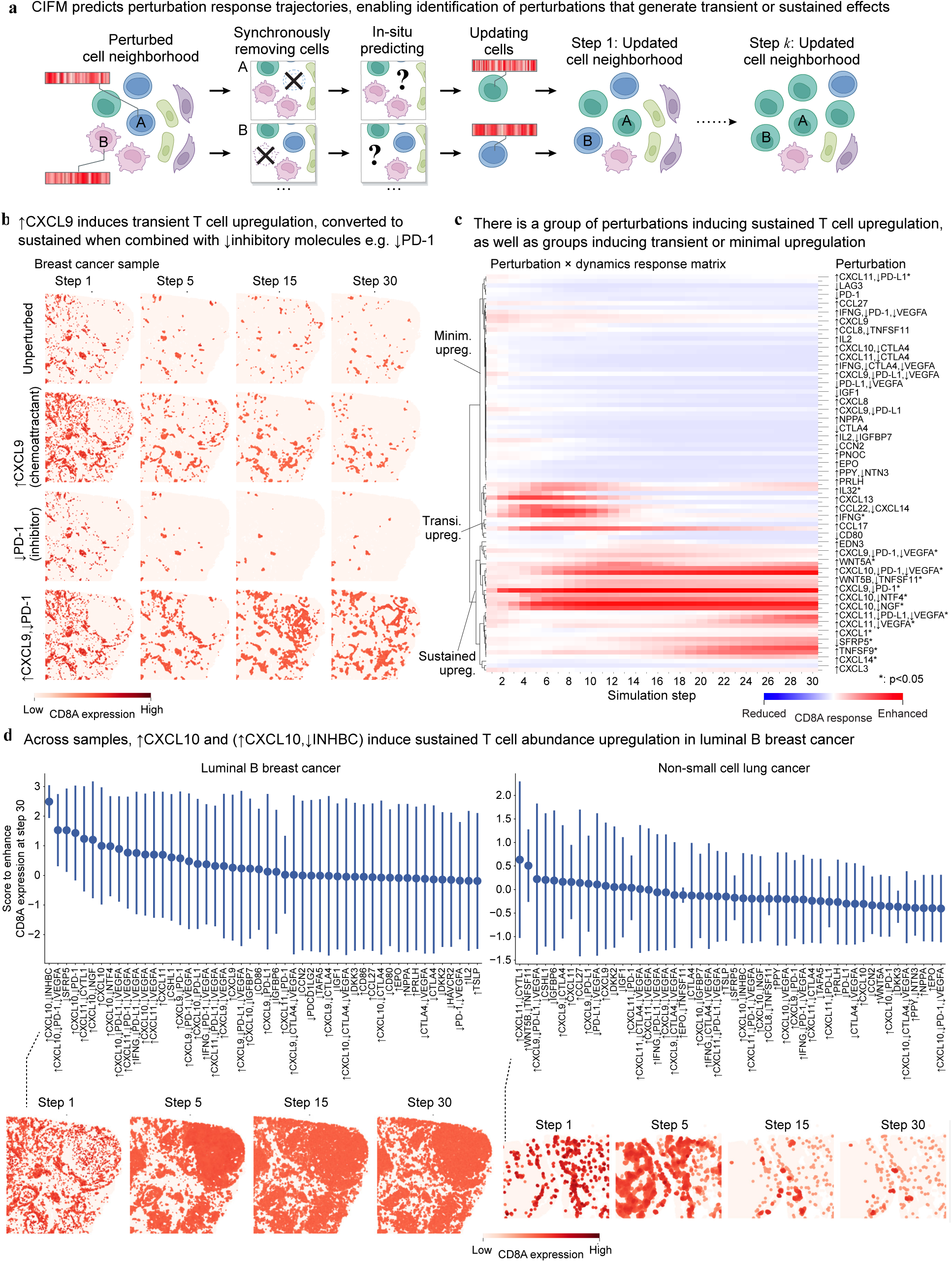
CIFM predicts perturbation response trajectories and separates transient from sustained effects. a, Autoregressive simulation. Cells in a perturbed neighborhood are synchronously removed, their transcriptional states predicted in situ, and the updated states returned to the tissue; iterating yields the tissue state at step k. b, CD8A expression in a breast cancer sample at simulation steps 1, 5, 15 and 30 under no perturbation, CXCL9 over-expression, PD-1 knockdown, and the combination. c, Perturbation-by-dynamics response matrix across 30 simulation steps, hierarchically clustered into sustained, transient and minimal up-regulation groups; asterisks denote *p <* 0.05. d, Perturbations ranked by predicted CD8A enhancement at step 30 in luminal B breast cancer (left) and non-small-cell lung cancer (right); points, mean; bars, s.d. across patients. Below, spatial CD8A response across simulation steps for the top-ranked perturbation in each tumor type.

## Methods

### Spatial transcriptomics data

Human spatial transcriptomics data were assembled from public resources (Table S1). Raw counts and spatial coordinates in micrometers were read into AnnData objects [60]; counts were normalized to 10^4^ transcripts per cell and log(1 + *x*)-transformed, with raw counts retained in a separate layer; and gene identifiers were harmonized to Ensembl across platforms.

Curation yields two nested corpora. The full curated corpus comprises **49,953,298 cells** across **281 tissue sections**, **39,551 genes**, six measurement platforms and fifteen anatomical sites. The model reported here was trained on the three-platform subset of that corpus, **22,951,580 cells** across **56 sections** and **18,289 genes**. Sections were partitioned into training, validation and test regions.

Candidate datasets are identified by an automated agent that surveys the literature and the public repositories continuously. The agent queries literature indices over rolling date windows for spatial genomics keywords, retrieves the full text of each hit through open-access and archive routes, and uses a language model to extract one structured record per dataset described in the paper, comprising the host repository, the accession, the measurement platform, species, tissue, disease state and whether the data are public and were newly generated. It queries the Gene Expression Omnibus and Zenodo directly over the same terms so that depositions not yet linked to a publication are also captured. The agent produces a ranked registry of candidates, currently 1,634 publications and 1,890 dataset records; records meeting the criteria above are downloaded, harmonised and appended to the corpus. The corpora reported here are a snapshot of that process.

**Table S1:** Sources of the curated corpus.

| Source | Cells | Platform(s) and provenance |
| --- | --- | --- |
| Base atlas | 33,091,583 | Xenium V1, Xenium Prime, Visium HD (10x Genomics public datasets) and MERFISH (Vizgen FFPE human immuno-oncology release; 18 sections spanning prostate, uterine, lung, colon, liver, breast, melanoma and ovarian tumours) |
| SPATCH[61] | 6,898,885 | CosMx, Stereo-seq, Visium HD (FF and FFPE) and Xenium; matched COAD, HCC and OV tumours |
| CosMx panels | 4,149,035 | CosMx; colon, frontal cortex, liver, lung, lymph node, pancreas |
| Visium HD new releases | 1,830,559 | Visium HD; breast, colon, kidney, lymph node, ovary ($\times 3$), pancreas, prostate |
| Colorectal MSI/MSS[62] | 1,388,060 | Xenium 5K; two colorectal sections |
| Skin MERFISH[63] | 1,201,886 | MERFISH; 114 regions (Zenodo 16795569) |
| Open-ST[64] | 1,097,769 | Open-ST; metastatic lymph node, 19 sections (GEO GSE251926) |
| Renal carcinoma[65] | 199,112 | CosMx; kidney (Zenodo 12730227) |
| Neurodegeneration[66] | 96,409 | Visium HD; Alzheimer’s disease, control, multiple sclerosis, glioblastoma, focal cortical dysplasia, Aicardi–Goutieres syndrome (Zenodo 16938034) |

### Model and training

CIFM is a geometric graph neural network that predicts the genome-wide transcriptional state of a cell from the states of the cells around it. Each tissue section is represented as a graph whose nodes are cells carrying their measured expression profile and spatial position and whose edges join spatially proximate cells. It is trained by self-supervision: within each cell neighborhood the transcriptome of one cell is masked and the network is trained to reconstruct it from its neighbors alone, using a decoder that predicts both which genes are detected and at what level, against a combined regression and classification objective. Training used 23 million cell microenvironments drawn from the corpus above. Models spanning 100 million to 4 billion parameters were trained under a common recipe to measure how accuracy depends on capacity, compared at a matched optimization budget; the model used for the simulations reported here is the 100-million-parameter configuration.

### Forward simulation of perturbation responses

A virtual perturbation sets the expression channel of one or more target genes to a fixed value in every cell of the tissue and holds it there for the duration of the simulation; combinatorial perturbations apply one such value per target. Tissue trajectories are generated by Monte-Carlo play-out: at each step every cell is masked in turn and its transcriptome resampled from its current neighborhood, all cells are updated simultaneously, and the updated tissue becomes the input to the next step, with cell positions and the graph held fixed throughout. Every simulation is run alongside a matched unperturbed simulation of the same tissue, and all reported responses are differences against that control at the same step. Two readouts are used: a single forward pass, which is what the large combinatorial screens report, and iterated play-out over thirty steps for the tumor trajectory analyses.

### Computational analyses

Prediction accuracy was assessed on held-out tissue regions against three baselines – averaging over neighboring cells, a spatially aware dimensionality reduction [48], and a random expression model – with the attainable ceiling defined by corrupting the measured data with simulated single-cell sequencing noise and passing it through the same evaluation. Predicted and measured profiles were annotated with a published cell-type classifier [49] and compared by the agreement of their ranked labels. Microenvironment representations were taken from the same forward pass with the central cell masked, visualized by UMAP, and used to predict tissue and disease state with whole sections held out of training. Tissue programs were defined and scored in later simulations. Perturbation screens were scored on marker genes and, for the trajectory analyses, grouped by hierarchical clustering of the response across steps. Over-expressed ligands were grouped by their annotated receptor using a public protein-interaction database [52], and the similarity of their genome-wide responses compared within and between groups.

### Experimental validation

T cell infiltration was measured in a transwell migration assay. BT-474 luminal B breast carcinoma cells were seeded into the lower chamber of a 96-well transwell plate with an 8 *µ*m pore membrane and allowed to attach; recombinant chemokines and blocking antibodies were added to the lower chamber; and human peripheral blood mononuclear cells, activated with a CD3/CD28 stimulus, were loaded into the upper insert. Chemokines were used at 100 ng mL*^−^*^1^ and blocking antibodies at 5 *µ*g mL*^−^*^1^. After 4 h at 37*^◦^*C the cells that had crossed into the lower chamber were collected, stained for CD3 and CD8, and counted by flow cytometry. Twenty-six conditions were assayed on a single plate with three to four replicate wells each, and counts are reported as a fold change relative to wells receiving tumour cells and mononuclear cells in medium alone. Replicates are wells within one experiment.

Simulated responses were additionally compared with two sets of previously published measurements. The first is organoid and xenograft data for NSD2 inhibition in prostate cancer [50]. The second is single-cell transcriptomes of physically interacting tumor and T cell pairs, obtained by cell–cell sequencing [51].

## Ethics

All human tissue data analyzed in this study are previously published, publicly available and deidentified, and were generated under the approvals and consents reported in the original studies cited in Table S1. No new human-subjects research was undertaken.

## Data and code availability

All spatial transcriptomics datasets analysed in this study are publicly available from the repositories and accessions listed in Table S1. Code is available from the corresponding authors on request.

